# Flexible modeling of large-scale neural network stimulation: electrical and optical extensions to The Virtual Electrode Recording Tool for EXtracellular Potentials (VERTEX)

**DOI:** 10.1101/2024.08.20.608687

**Authors:** Anne F. Pierce, Larry Shupe, Eberhard Fetz, Azadeh Yazdan-Shahmorad

## Abstract

**Background:** Computational models that predict effects of neural stimulation can be used as a preliminary tool to inform *in-vivo* research, reducing the costs, time, and ethical considerations involved. However, current models do not support the diverse neural stimulation techniques used *in-vivo*, including the expanding selection of electrodes, stimulation modalities, and stimulation paradigms.

**New Method:** To develop a more comprehensive software, we created several extensions to The Virtual Electrode Recording Tool for EXtracellular Potentials (VERTEX), the MATLAB-based neural stimulation tool from Newcastle University. VERTEX simulates input currents in a large population of multi-compartment neurons within a small cortical slice to model electric field stimulation, while recording local field potentials (LFPs) and spiking activity. Our extensions to its existing electric field stimulation framework include allowing multiple pairs of parametrically defined electrodes and biphasic, bipolar stimulation delivered at programmable delays. To support the growing use of optogenetic approaches for targeted neural stimulation, we introduced a feature that models optogenetic stimulation through an additional VERTEX input function that converts irradiance to currents at optogenetically responsive neurons. Finally, we added extensions to allow complex stimulation protocols including paired-pulse, spatiotemporal patterned, and closed-loop stimulation.

**Results:** We demonstrated our novel features using VERTEX’s built-in functionalities, with results in alignment with other models and experimental work.

**Conclusions:** Our extensions provide an all in one platform to efficiently and systematically test diverse, targeted, and individualized stimulation patterns.

## 1. Introduction

Neural stimulation has significant history and promise for treating neurological disorders characterized by damaged or aberrant neural activity, such as movement disorders, epilepsy, and stroke. However, the effectiveness of stimulation-based treatments has variable outcomes across clinical and preclinical trials. This inconsistency is attributed to the use of non-individualized stimulation and diverse methods employed across experiments, including variations in electrode types, spatial and temporal stimulation dynamics, and open versus closed-loop approaches. While it is critical to investigate methods that consistently yield optimal outcomes, *in-vivo* experiments are time-intensive, expensive, and raise ethical considerations regarding the use of humans and animals. Consequently, before conducting *in-vivo* experiments, computational modeling can be used as a fast and cost-effective method to predict effects of stimulation under various conditions. The results could inform and reduce the number of subsequent *in-vivo* experiments, and aid in the development of reliable, individualized, and targeted therapeutic treatments.

While existing software can predict effects of neural stimulation, most models simulate neural activity in large populations of neurons lacking realistic biophysical properties, or simulate activity in only a few neurons that possess complex, neurophysiological characteristics. Since *in-vivo* neural stimulation induces both local and network-wide effects that contribute to its therapeutic outcomes, it is crucial to have a model suited for an extensive network of neurons while maintaining realistic properties. The Virtual Electrode Recording Tool for EXtracellular Potentials (VERTEX) is a MATLAB-based software designed to simulate local field potentials (LFPs) and spike timing in response to electrical stimulation in a large population of neurons within a multi-layer slice of cortex (Tomsett *et al* 2015; Thornton *et al* 2019). VERTEX simulates neuron types, compartments, densities, and connectivity properties based on empirical research, which lends to realistic neuron characteristics. Additionally, VERTEX generates network dynamics using imported spike times or using the adaptive exponential integrate and fire (AdEx) model (Brette 2005), which can mimic the firing patterns of many different neuronal cell types. Together these features achieve a balance between complexity and practicality to give rise to realistic spiking patterns and LFP calculations, making VERTEX uniquely suited to efficiently test the effects of electrical stimulation-based approaches in a slice of cortex prior to *in-vivo* experiments.

However, VERTEX has constraints that hinder its ability to model the wide range of approaches used *in-vivo*. These include a restrictive and cumbersome electrode design process, a suboptimal electrical stimulation waveform, a single stimulation modality, and few stimulation protocols. To overcome these limitations, we developed several extensions to VERTEX to broaden its capabilities. We first developed a new script that enables electrical stimulation with biphasic waveforms and facilitates rapid creation and modification of electrode shape, number, and positioning. Next, we created a model to simulate optogenetic stimulation by converting irradiance to input current. This represents a significant advancement since optogenetics has become a highly prevalent method to deliver targeted stimulation. Finally, we added the capability to deliver three stimulation protocols including paired-pulse stimulation, spatiotemporal patterned stimulation, and closed-loop stimulation. We demonstrate our novel extensions using VERTEX’s built-in spiking and LFP recordings. These novel features allow users to test a vast array of stimulation approaches, providing a highly adaptable LFP simulation tool.

## 2. Materials and Methods

All simulations used the following settings unless otherwise noted. The size of the simulated tissue block was 1.5 × 1.5 × 2.6 mm deep with virtual electrodes for LFP recording sites spaced in a 3 × 3 × 6 grid to capture activity in each cortical layer. We used the 15 neuron-group VERTEX model developed by Tomsett *et al* (2015), which incorporates the biophysical and connectivity patterns of 15 distinct types of cortical neurons, each characterized by unique features such as compartmental structure, soma location, projecting layer, firing rate, number of synapses and synapse dynamics. Our resulting networks contained approximately 224 thousand neuron units and 569 million connections. VERTEX calculates LFPs by summing the membrane potentials of each compartment, weighted by their distance from the virtual electrodes. Neuronal spiking was driven by synaptic currents as well as stochastic AdEx input currents. The means and standard deviations for the AdEx input currents used in Tomsett *et al* (2015) result in large gamma oscillations that can mask other evoked potentials. To reduce the model’s inherent gamma oscillations to levels low enough to not obscure stimulus-evoked LFPs, we chose to scale the mean and standard deviations of the AdEx input currents by 1.125x and 1.75x. Simulations were run remotely on the Neuroscience Gateway (Sivagnanam *et al* 2013) computer cluster or on a local PC (AMD 7800X3D CPU with 128GB memory) and generally required about 1 hour run time per 1 second of simulated time to complete. A list of added or modified code modules are reported in Supplementary Table 1.

**Table 1.** Parameters for estimating irradiance at coordinate (x, y, z) in millimeters.

| Parameter | Blue light (473 nm) | Amber light (594 nm) |
| --- | --- | --- |
| $\tau$ | 0.39 mm | 0.38 mm |
| $a$ | 92 | 8.8 |
| $h$ | 1.14 | 1.67 |

### 2.1. Electric field stimulation: parametric electrodes and biphasic stimulation

VERTEX has built in support for electric field stimulation with demonstration code for monophasic stimulation through a single pair of differential electrodes positioned horizontally through the model tissue slice. The 3D electrode topology is created in an external 3D modeling application and imported into MATLAB. The reliance on separate software requiring multiple cumbersome steps limits rapid modification and parameterization of electrodes. To overcome this limitation, we implemented a new script for electric field stimulation that removes the dependence on an external 3D application. A function called within this script parametrically creates electrode topologies directly in MATLAB, allowing easy modification of the electrode shape, the number of electrode pairs, and the positioning of the electrodes within the tissue volume. This function generates the same format of tessellated 3D geometry that VERTEX would otherwise import from an external application and that is used by MATLAB’s PDE Toolbox to build a finite element model of the electric potential in the tissue volume resulting from a potential difference between the electrodes (Thornton *et al* 2019). We demonstrate the benefit and versatility of this user-friendly feature with single and multiple pairs of tapered tip and surface patch electrodes oriented perpendicular to the ventral surface of the modeled tissue, resembling electrodes in the Utah Array or an Electrocorticography (ECoG) array (Fig 1).

**Figure 1.**
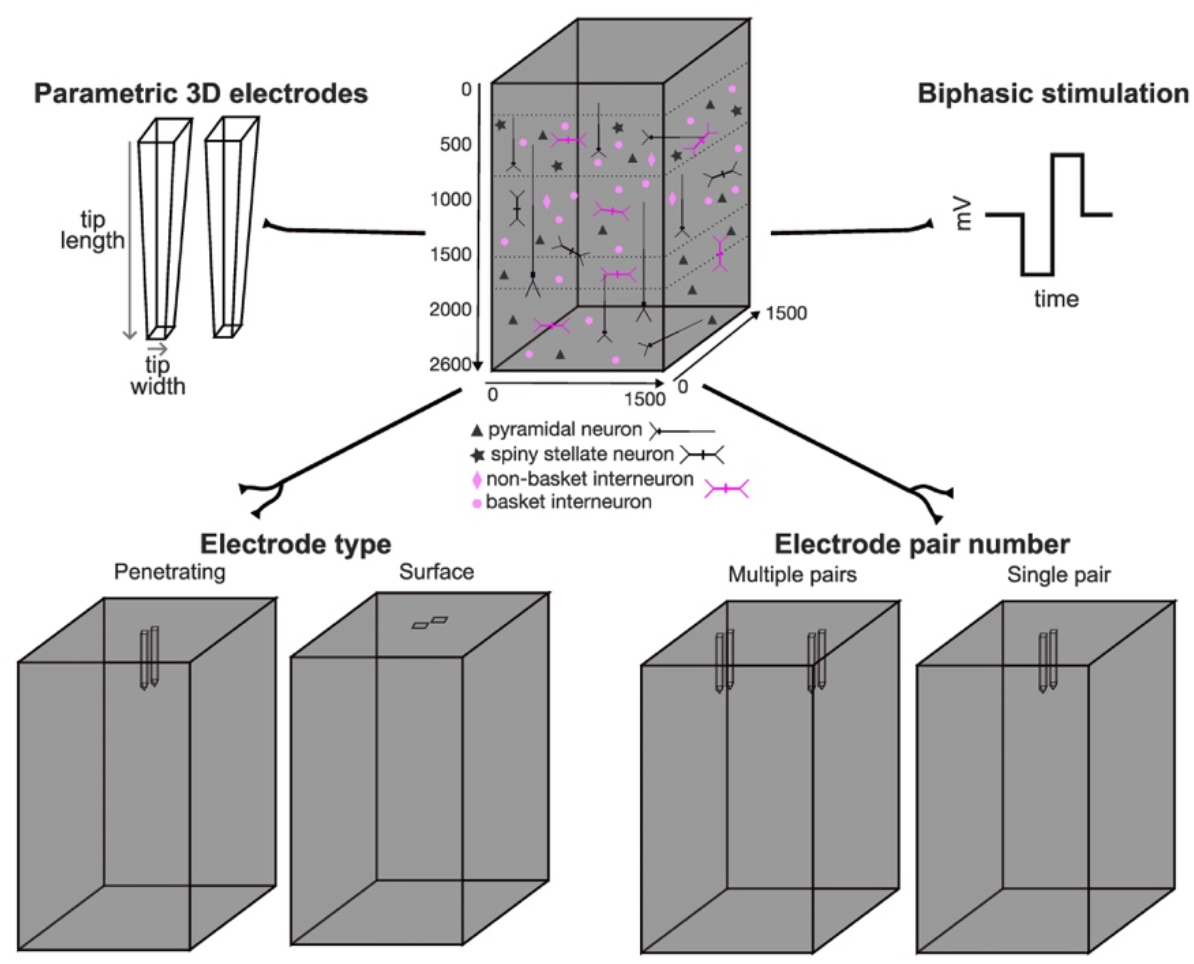
Added features for electric field stimulation. VERTEX defines a tissue volume where a variety of modeled neuron types are placed. We introduce several features to increase flexibility and versatility when defining electrode and stimulation parameters in the tissue volume. Electrode positions, lengths, and widths are parametrically defined using the MATLAB PDE toolbox. The electrode geometry can represent penetrating or surface electrodes in a single pair or multiple pair configuration. Biphasic stimulation is modeled by inverting the electric field halfway through the pulse duration.

Additionally, in this script we introduce features that significantly expand the range of stimulation options. For example, we add the ability to modify stimulation timing and pulse parameters during an ongoing simulation, a feature particularly beneficial for closed-loop stimulation.

Lastly, rather than restricting stimulation to a single pair of differential electrodes with monophasic waveforms, our code accommodates multiple pairs of stimulating electrodes that allow biphasic, bipolar stimulation. This stimulation waveform is more commonly used in clinical settings, as it is less likely to cause abnormal neuronal activity, tissue damage, and electrode degradation compared to monophasic stimulation (Yuan *et al* 2021). We perform biphasic stimulation by inverting the electric field halfway through the stimulus duration. While this is constant voltage stimulation, the VERTEX tissue model is purely resistive and the current applied can be estimated from the tissue conductivity, electrode surface area, and the electric field calculated by the Matlab Partial Differential Equation (PDE) Toolbox. These novel features broaden the range of electrode and stimulation settings available, facilitating comprehensive investigations into effective parameters for modulating neural activity.

### 2.2. Modeling optogenetic stimulation

Optogenetics has become a commonly used technique to rapidly modulate neural activity in neurons expressing exogenous light-sensitive ion channels. By applying light to the targeted region, the light-sensitive ion-channels open and induce a photocurrent in the affected cells. We created a novel script to model optogenetic stimulation using VERTEX’s built-in functionality for adding input currents to neuron units. These currents can vary with time and may be turned on and off to model photocurrents.

Light-sensitive units are defined in the script, allowing users to specify which cell types or layers to set as light-responsive. The light source for optogenetic stimulation is typically a laser which projects light of a specific wavelength through an optical fiber. The laser’s radiant power (P in mW) and the fiber’s radius (r in mm) are additional user-defined parameters in the script, and control the intensity of the stimulation with the initial irradiance (E_0_) at the tissue surface beneath the optic fiber defined by Equation 1.

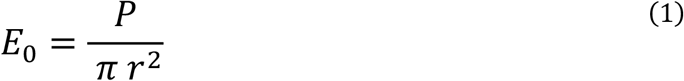

Irradiance at depth (z) directly below the light source is modeled by fitting both an exponential and geometric decay to data from Yizhar *et al* (2011), where an optical fiber was lowered through a tissue block to measure light transmission through unfixed brain tissue. The 10% and 1% light transmission contours provided the percentage of light remaining at depth and lateral distance from the optical fiber center line. The depth (z) of these contours is measured for both 473 nm and 594 nm light and fit to Equation 2. The ratio (h) of depth to half-width (at half-depth) of the 1% contours is used to calculate a scaled lateral distance (l) to create a 3-dimensional estimate of irradiance at any (x, y, z) coordinate offset from the light source tip. Parameter fitting values are shown in Table 1 and the irradiance estimate (mW/mm^2^) is shown in Equation 3. When optical stimulation is initiated, irradiance values for each light source are calculated for each light-sensitive unit at its soma position.

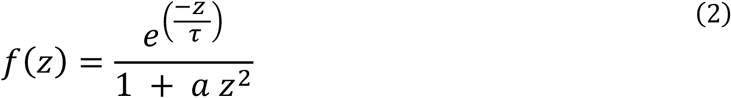

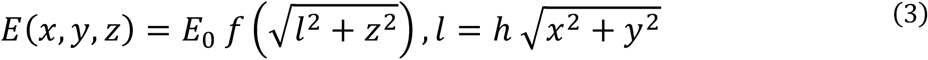

There are several theoretical models for converting irradiance to photocurrents for various opsins. We chose Foutz *et al* (2011) for modeling Channelrhodopsin-2 (ChR2) with 473 nm light because it demonstrated photocurrent responses across various neuronal compartments and irradiance levels. For Chronos and vfChrimson we selected models that offered a comprehensive investigation of photocurrent dynamics across several parameters - including pulse width and frequency, irradiance, and light wavelength - with results congruent with experimental work (Saran *et al* 2018; Gupta *et al* 2018). For Jaws, we relied on work from Chuong *et al* (2014) as it provided experimental results for photocurrents elicited by various irradiance values. Peak photocurrent estimates (in picoamps) for irradiance levels E (in mW/mm^2^) were fit with Equation 4 for ChR2, Equation 5 for Chronos, Equation 6 for vfChrimson, and Equation 7 for Jaws.

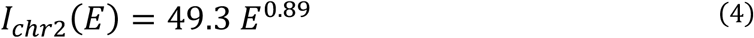

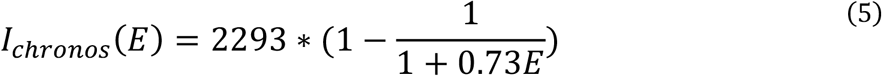

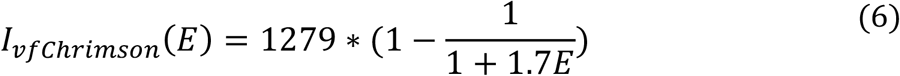

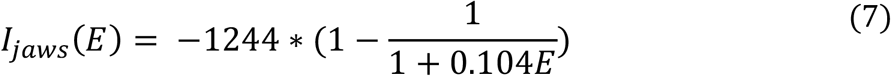

Photocurrent dynamics are written into a VERTEX input model that handles optogenetic stimulation. This is a step function with exponential on and off dynamics to simulate the rise and fall of an input current to a precalculated value during the application of a light pulse. The time-constants used for the on and off mechanics for ChR2 are τ_on_ = 1.5 ms, τ_off_ = 11.6 ms (Mattis *et al* 2012), for Chronos are τ_on_ = 0.65 ms, τ_off_ = 3.6 ms (Saran *et al* 2018,) for vfChrimson are τ_on_ = 1.0 ms, τ_off_ = 2.7 ms (Gupta *et al* 2019), and for Jaws are τ_on_ = 3.6 ms, τ_off_ = 4.2 ms (Chuong *et al* 2014).

### 2.3. Stimulation paradigms

We created new scripts for three stimulation paradigms: paired-pulse, spatiotemporal patterned, and closed-loop stimulation. Each paradigm can use either electrical or optogenetic stimulation. Paired-pulse and spatiotemporal patterned stimulation both involve delivering stimulation at multiple sites with temporal delays between them. These spatial and temporal properties can induce spike-timing-dependent plasticity (STDP), a biological phenomenon based on spike timing differences between the postsynaptic unit (firing at time t2) and the presynaptic unit (firing at t1) with the spike-timing difference defined as Δt = t2 – t1. Positive differences strengthen while negative differences weaken connectivity between the pre- and postsynaptic unit. STDP is built into VERTEX synapse models to allow changes in connection strengths between units. In this STDP implementation, each time the pre- or postsynaptic neuron fires, there is an update to synapse connectivity, where two exponential functions (per synapse), each with unique decay times for the pre- and postsynaptic neuron, dictate the degree of synaptic connectivity change.

Although VERTEX demonstrates a form of paired-pulse stimulation with STDP, it currently only supports paired-pulse stimulation using a single pair of electrodes at the same site, whereas paired-pulse stimulation is typically administered at separate sites. Since this paradigm does not represent the typical protocol used *in-vivo*, we created a novel script for paired-pulse stimulation where stimulation is applied at distinct sites. Additionally, we created a new script to deliver spatiotemporal patterned stimulation, where stimulation can be applied to a greater number of sites with varying amplitudes and pulse delays between sites.

The third paradigm we support is closed-loop stimulation, where stimulation is delivered in response to on-going activity. In biophysical experiments, stimulation can be administered in response to behaviors, neural activity such as LFPs or single unit activity, and peripheral activity including signals from electromyography. In VERTEX, closed-loop stimulation is largely limited to recorded LFPs and spike times. We have implemented two forms of closed-loop stimulation, both of which are dependent on LFP measurements. The first is cycle-triggered stimulation where a stimulus pulse is delivered based on the amplitude and phase of the filtered LFP recorded on a single recording electrode. The second closed-loop paradigm is amplitude-adjusted stimulation where the amplitude of stimulation is adjusted to keep the magnitude of an LFP channel within a certain range. Both methods require transferring partial LFP values between the parallel MATLAB processes used to accelerate VERTEX so that each process has a complete copy of the LFP at each recording site.

## 3. Results

### 3.1. Optogenetic stimulation

To get an estimate of light penetration through the modeled tissue, we generated contour plots of irradiance at depth and lateral distance for 473 nm and 594 nm light using a single light source with a radius of 100 µm (Fig 2A). Both contour plots have a dramatic drop-off in irradiance in the modeled tissue, though the fall-off of 594 nm irradiance is more gradual compared to 473 nm. The depth of light penetration shown here for 473 nm and 594 nm light is consistent with previous *in-vivo* work, reflecting greater tissue penetration of longer wavelengths due to reduced light absorption and scattering (Senova *et al* 2017). Our models for converting irradiance to current are demonstrated in Figure 2B. For each of the four modeled opsins - ChR2, Chronos, vfChrimson, and Jaws - we show current induced by a 5ms light pulse across several irradiance values. In accordance with biophysical experiments, we found Chronos to have high sensitivity at low irradiance values (Klapoetke *et al* 2014). To highlight the diverse effects of optogenetic stimulation on spiking activity and LFP generation across the different opsins, we show simulations for each opsin under identical stimulation parameters (Fig 2C). Each simulation displays the average spiking and LFP response following a 5ms light pulse, where all units were set as light-responsive, averaged across 100 pulses. Each simulation used a light power of 7.2 mW and radius of 100 µm. To calculate tissue maps of spike-rate changes evoked by stimulation, unit spike times were divided spatially into 25 µm bins based on soma positions within the tissue volume. Baseline spiking rates were calculated for each bin by summing spike counts along either the Z axis (top-down view) or Y axis (tissue side-view) for the 50 ms time-window preceding stimulus onset times. Spike-rate responses were similarly calculated for the 5 ms stimulus duration. Percent increases in spiking were plotted on log scales to highlight smaller changes. For each simulation, these maps are shown from top-down (top row) and side-view perspectives (middle row), along with a stimulus-triggered-average (STA) of the surface LFP (bottom row).

**Figure 2.**
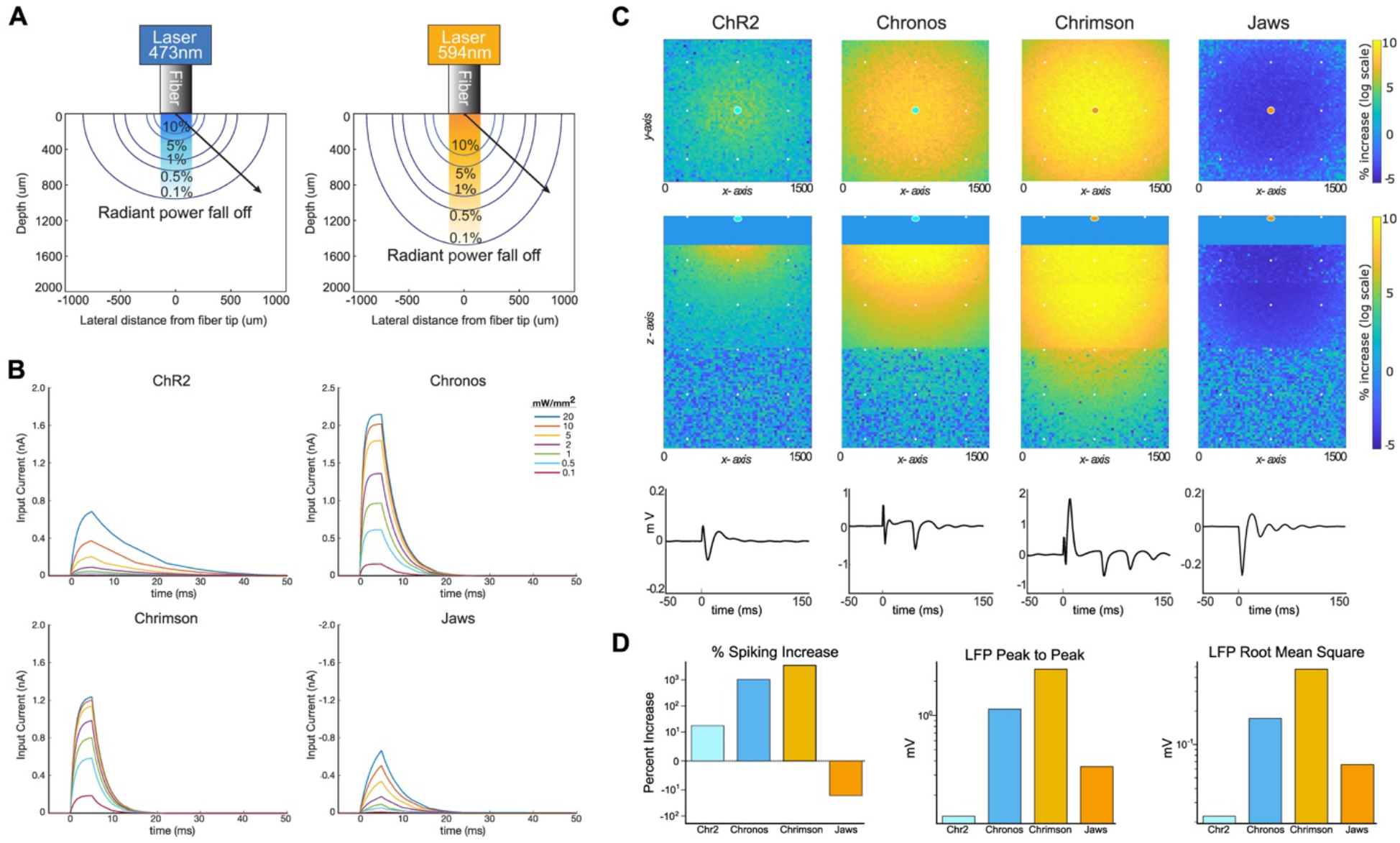
Modeling optogenetic stimulation. A) Light spread through tissue is modeled for blue light (473 nm) and amber light (594 nm) to determine photocurrent responses at different irradiance values. Radiant power fall off is due to both light absorption and geometric fall-off with distance. B) Photocurrent rise and decay are modeled for each of the four opsins using distinct exponential functions and are shown here in response to a 5 ms light pulse across several irradiance values. C-D) A simulation was run for each of the four opsins where 100 stimuli events were delivered. Each stimuli event consisted of a 5ms pulse using the same optogenetic parameters (fiber radii = 100µm; light power = 7.2mW; all neuron unit groups set as light-responsive). The simulation modeling the ChR2 and Chronos used the 473 nm/blue light model whereas the simulations modeling Chrimson and Jaws use the 594 nm/amber light model. C) The top-down (top row) and side-view (middle row) through the tissue show the percent change in spiking activity on log scales during the 5 ms of stimulation compared to a baseline period (50 ms prior to stimulation). The bottom row demonstrates LFPs at the surface-center recording electrode. D) Three measures of stimulus response strength shown in 2D: Percent increase in spiking during the stimulus (top), LFP peak to peak (middle), and root mean square (bottom) of the average surface LFP response during the 100 ms following the stimulus.

We quantified the stimulus response strength across simulations using 3 measures - percent spiking increase, LFP peak to peak, and the LFP root mean square (Fig 2D). Although ChR2 and Chronos simulations both used the blue-light model, Chronos stimulation evoked significantly greater spiking and LFP response than ChR2 stimulation, which can be attributable to Chronos’ increased light sensitivity and faster kinetics. While vfChrimson has lower light-sensitivity than Chronos, we found that vfChrimson activation led to increased spiking, spatial spread of spiking, and LFP response. This enhanced response is likely due to the use of amber light for vfChrimson activation, which penetrates tissue more deeply compared to blue light activation used in the Chronos simulation. As expected for an inhibitory opsin, the Jaws stimulation resulted in decreased spiking activity and a negative change in LFP response.

In addition to selecting which opsin to simulate, users can customize various stimulation parameters, including the light power, light radius, and which neuron unit groups to designate as light-responsive. Figure S1 demonstrates how modifying these parameters can influence spiking activity and LFP responses, with effects that range from subtle to pronounced. When examining light settings, we found that increasing the initial light power or decreasing the light radius, while maintaining the same light power, led to greater spiking activity, spatial spread of spiking, and LFP responses (Fig S1A-B). Additionally, in Figure S1D-E, we validated the ability of our model to allow cell type specific stimulation. When comparing vfChrimson activation in all units, excitatory units only, and inhibitory units only, we observed that excitatory units were primarily driving the maximum LFP response, whereas inhibitory units were regulating post-stimulation oscillations.

### 3.2. Paired-pulse stimulation with spike-timing-dependent plasticity

In Figure 3, we demonstrate our paired-pulse stimulation paradigm, combined with several of our extensions to electric field stimulation, including biphasic stimulation at multiple electrode pairs with a programmed delay. In this simulation we enabled VERTEX’s built in STDP feature that requires using a script where defined STDP parameters govern the temporal dynamics and degree of connectivity change. Based on work shown in Bi and Poo (2001), we set the decay time constants for the exponential curves to 17ms and 34ms for positive and negative Δt, respectively such that small values of Δt give the largest changes and large values of Δt give exponentially smaller changes (Fig 3A). The amplitude for the weakening function was set at 0.53 times that of the curve for the strengthening function to provide slightly more area under the weakening curve. This helps prevent run-away connection strengths from random activity since there is no homeostasis function. The maximum change can be modified but is normally set between 0.001 and 0.005 nanosiemens (nS). Connection magnitudes can be limited and are normally restricted to the range between 0.001 and 4.0 nS.

**Figure 3.**
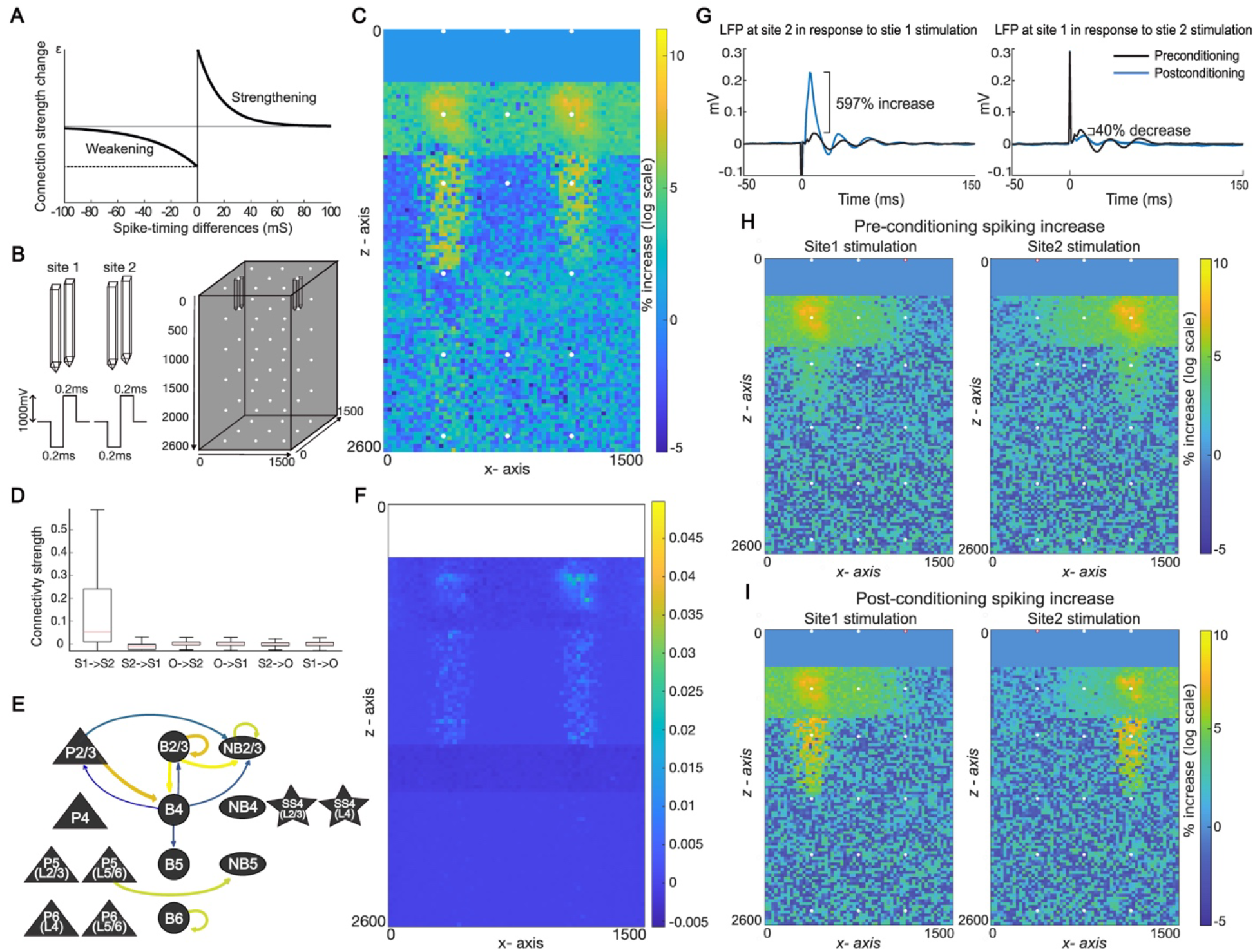
Paired-pulse conditioning. A) Schematic of STDP principle. B) Schematic of paired-pulse electric field stimulation and placement of stimulating (black outline) and recording electrodes (white dots) in the tissue slice. C) Side-view of percent increase in spiking activity in log scale during the 10 ms window after the stimulus onset. D) Mean connection strengths from different sites. Mean connection strength from Site1 to Site2 increased while that from Site2 to Site1 decreased. Mean connection strengths involving units outside (“O”) one or both sites remained largely unchanged. E) Neuron groups (arranged vertically by cortical layer) with only the largest mean connection-strength changes shown. Size and color of arrows reflect degree of change with blue to yellow reflecting increasing strength changes. “P” = pyramidal neuron, “B” = basket interneuron, “NB” = non-basket interneuron, “SS” = spiny stellate neuron. Layer abbreviations within parentheses represent the projection layer. F) Side-view of connection strength changes showing that the largest changes occur to units within 100 um radius of the site’s (x,y) locations in both layer 2/3 and layer 4. G) Stimulus evoked LFP responses for electrodes in layer 2/3 near site 2 or site 1 for both the preconditioned and post-conditioned network. H) Increased spiking activity in the 5 ms window after test stimulation for each site in the preconditioned and I) post-conditioned networks.

In Figure 3 we used the original AdEx input-current scalers since plasticity reduces the network’s inherent oscillations to levels low enough to not obscure the stimulus-evoked responses. This also allowed for larger stimulus responses in deeper layers which resulted in brief oscillatory activity that dampened out within 100 ms. Network connection strengths were initialized from the results of running a non-stimulating network for 30 seconds with STDP turned on, allowing the paired-pulse conditioning to begin with a more stable distribution of connection strengths and very low LFP oscillations.

Paired-pulse conditioning was simulated using electric field stimulation at two sites separated by 750 µm in the middle of layer 2/3 (Fig 3B). The electrode tips were modeled after a commonly used microelectrode array and used 50 µm tip lengths and 35 µm base diameters. The bipolar tips were placed 100 microns apart. 100 paired stimulation events were delivered where stimulation at the second site was delayed 5 ms from the first. 1000 mV biphasic-bipolar stimulation was delivered to each site in brief 0.4 ms pulses (0.2 ms each phase). This produced an estimated constant current stimulation of 65 µA at each site since the VERTEX tissue model is purely resistive.

Stimulus times were used to calculate post-stimulus changes in spiking activity and create STAs of resulting LFPs similar to graphs for optogenetic stimulation in Figure 2 and Figure S1. To capture effects at both sites in Figure 3C, spike-rate responses were calculated for 0-10 ms after stimulation was delivered at the first site. Network connection strengths were saved before and after paired-pulse conditioning. Figure 3D-F shows connection strength changes (post – pre) by stimulation site, unit type, and unit location. Together, these results indicate an increase in connection strength from Site1 to Site2, and a slight decrease from Site2 to Site1. We used the network connection strengths before and after paired-pulse conditioning and compared the response to a single pulse stimulation at Site 1 or Site 2 (Fig 3G-I). After conditioning there was a 597% increase in the LFP peak value at Site 2 in response to Site 1 stimulation, and a 40% decrease in LFP peak value at Site 1 in response to Site 2 stimulation (Fig 3G). Some of the LFP changes were due to the increased spiking activity (post-conditioning) in layer 4, which was largely symmetrical for both Site1 and Site2 (Fig 3I).

### 3.3. Spatiotemporal patterned stimulation

In Figure 4 we illustrate a simulation using our extensions for spatiotemporal patterned and optogenetic stimulation. Four optogenetic stimulation sites are placed in each of the four surface quadrants of the tissue slice: lower-left, upper-right, lower-right, and upper-left (Fig 4A). These sites were stimulated, in that order, by 5 ms light pulses, each separated by 15 ms between the start of each light pulse. We used the ChR2/473 nm light model with light sources of 100 µm radius. The initial light power was 7.2 mW for each light source and all neuron groups were set as optogenetically responsive (Fig 4B). This train of pulses was repeated every 200 ms for 20 seconds. Stimulus triggered spiking activity and LFP averages were calculated as before. Figure 4A shows the stimulation response centered at each site with refractory responses visible for previous stimulation sites. Spiking activity after the fourth stimulation site is shown from the side-view (Fig 4C) and top-down view for individual layers (Fig 4D). Graphs aggregating activity within individual layers show localized spiking activity during the stimulus to layer 2/3 and 4 with lingering refractory responses from the previous site in layers 4 and 5. This aligns with experimental work showing that stimulation at the cortical surface reduced firing rates in deeper cortical layers (Yazdan-Shahmorad *et al* 2011; Yazdan-Shahmorad *et al* 2013). The STAs of the LFP show evoked potentials for each light pulse that do not completely decay before the next light pulse is delivered (Fig 4E).

**Figure 4.**
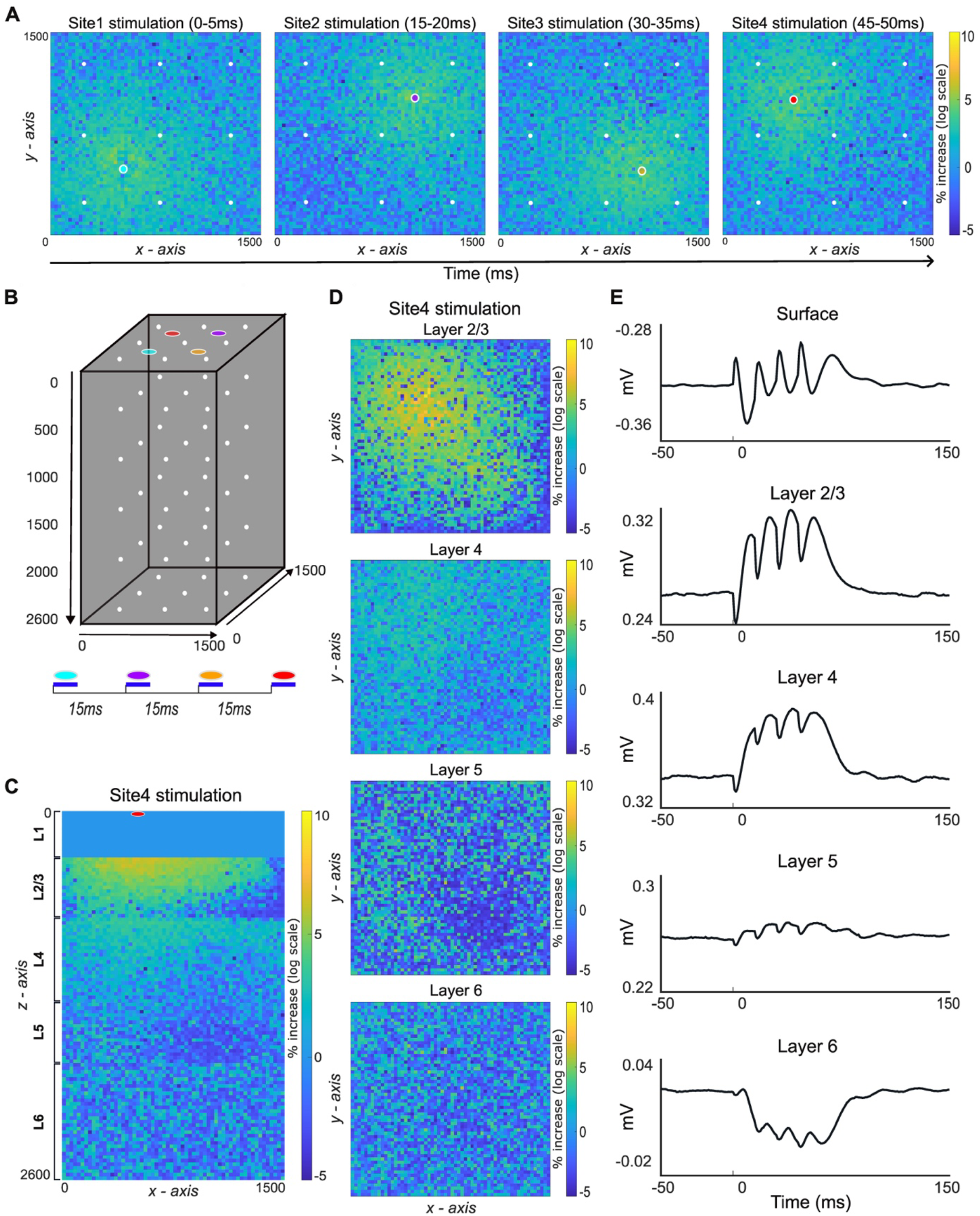
Spatiotemporal patterned stimulation. A) Top-down view of increased spiking activity on log scales in response to four consecutive optical pulses delivered at 15 ms intervals to different sites in the tissue slice. B) Placement of each of the light sources (colored dots) and recording electrodes (white dots) in the tissue slice. Timing of stimulation for each light source shown on bottom. Dark blue bars indicate 5ms light pulse durations. C) Side-view of increased spiking activity from the fourth stimulus site. D) Top-down view of spiking activity aggregated by layer after stimulation at the fourth site. E) LFP averages for the center column of recording electrodes aligned at the first pulse.

### 3.4. Cycle-triggered closed-loop stimulation

Figure 5 shows our cycle-triggered closed-loop stimulation using our new features to electric field stimulation to deliver biphasic stimulation at a single pair of differential electrodes in layer 2/3 (Fig 5A). To remove baseline signal-shift and reduce high-frequency noise, a 20-30 Hz band-pass filter was applied to the surface recording electrode located above the stimulating electrode. Stimulation was triggered by a rising filtered LFP with a magnitude threshold of 5 µV and a refractory period of 100ms (Fig 5B). Similar to the paired-pulse conditioning in Figure 3, 1000 mV biphasic-bipolar stimulation was delivered in brief 0.4 ms pulses. The pre-stimulus oscillation appears in the LFP STA with the average evoked LFP response (Fig 5C). The simulation ran for 30 seconds, with stimulation applied only between 5-25 seconds of simulation time.

**Figure 5.**
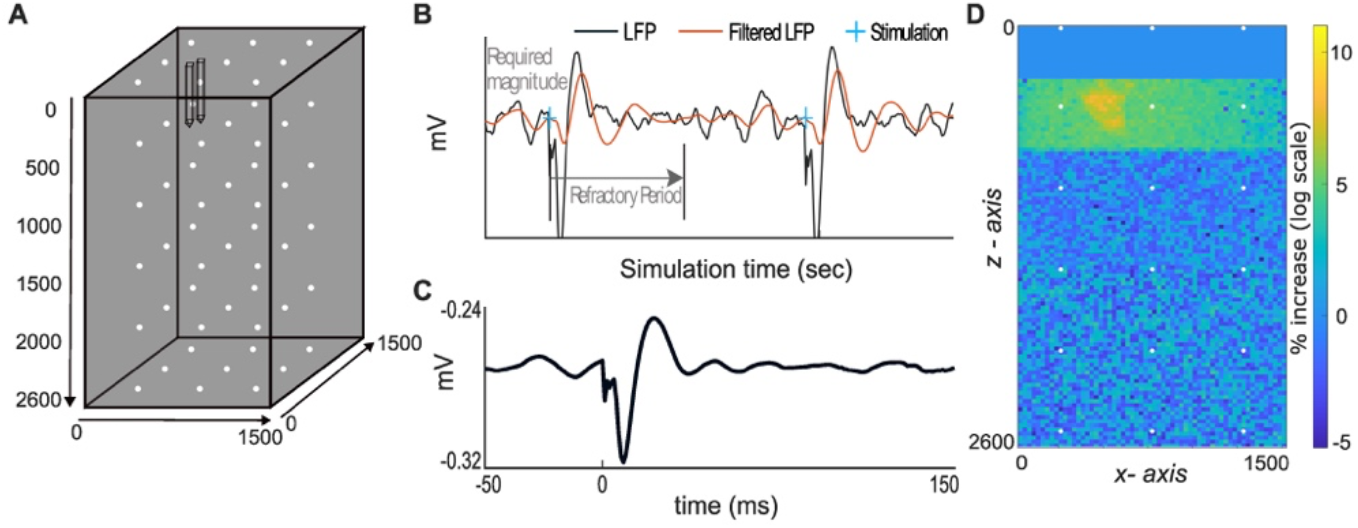
Closed-loop stimulation. A) Placement of stimulating (back outline) and recording electrodes (white dots) in the tissue slice. B) Schematic of stimulation triggered by rising LFP. C) STA of the unfiltered LFP. D) Side-view of the change in spiking activity (0-10 ms post-stimulus) in log scale after stimulation.

Within these 20 seconds, the filtered LFP met criteria to trigger stimulation 47 times. The location of increased post-stimulus spiking-activity is centered on the stimulation site with activity spreading primarily through layer 2/3 (Fig 5D).

## 4. Discussion and Conclusions

### 4.1. Novel extensions

We present novel extensions for VERTEX that enhance the software’s ability to model a diverse range of *in-vivo* stimulation approaches. First, we introduce a script that adds several new features for electric field stimulation, including the ability to parametrically create 3D electrodes using built-in MATLAB functions. This eliminates the need for external 3D modeling software, allowing users to easily create and position different electrode shapes, such as patches on the cortical surface or tapered electrodes penetrating the tissue. Additionally, we implemented the ability to deliver biphasic instead of monophasic stimulation, a stimulation waveform commonly used clinically due to more precise spatial targeting, tissue safety, and electrode longevity. Finally, we enable stimulation with programmable delays, which can be used to deliver stimulation with complex temporal and spatial patterns that can induce synaptic plasticity. These added functionalities facilitate users to easily test various electrode types and stimulation settings to identify the approaches that produce results most similar to their targeted outcomes.

Another key feature we implemented is the ability to model optogenetic stimulation. Over the past twenty years, optogenetics has become a widely adopted neuroscience technique used to control neural activity with spatial and temporal precision (Deisseroth 2015). While optogenetics is primarily used in preclinical research, experimentalists are beginning to adapt optogenetics for clinical trials (Gao *et al* 2023). Our extension offers extensive parametrization, developed specifically to mimic the technical choices available to experimentalists. For example, we model optogenetic stimulation with four popular opsins -Channelrhodopsin2, Chronos, vfChrimson, and Jaws - each having their own biophysical advantages and limitations. For instance, longer wavelengths of light, such as 594 nm used for vfChrimson and Jaws for neuronal activation and inhibition, respectively, can penetrate the brain deeper than 473 nm used for ChR2 and Chronos activation. Equally important, a user may require tightly regulated temporal stimulation, making the fast on/off kinetics of the Chronos model desirable. Another method commonly employed *in-vivo* is to select an opsin with a promoter that targets specific cell types. We support this technical approach by allowing specification of which neuron groups are light-responsive, thereby enabling stimulation of specific cell types or layers. Thus, depending on the desired depth of stimulation, kinetics of each opsin, neuronal target, and available resources, users can modify variables that best meet their needs. To our knowledge, our extensions provide the most comprehensive tool to model network wide effects of optogenetic stimulation under diverse parameters.

Finally, we developed open and closed-loop stimulation protocols that permit users to model stimulation with versatile temporal and spatial properties. Each protocol can be used with electric field or optogenetic stimulation. Furthermore, though we only demonstrate STDP with paired-pulse stimulation, STDP can be enabled for each protocol. Simulations with STPD take much longer to run due to the extra overhead and calculations (e.g. paired-pulse stimulation with STDP takes 3 times longer to run than paired-pulse stimulation without STDP enabled), but they can provide information on how connection strengths could change under specific interventions. For instance, compared to pre-conditioning, after paired-pulse conditioning, we found that stimulation delivered at Site1 resulted in a 597% larger LFP response at Site2 (Fig 3). These results are similar to other population-based neuron simulation tools. For example, the integrate-and-fire model developed by Shupe and Fetz (2021) found a 600% increase in evoked response after delivering paired pulse stimulation using a similar delay. More important, these results are congruent with *in-vivo* work showing that paired-pulse stimulation can strengthen connectivity between stimulation sites (Yazdan-Shahmorad *et al* 2018; Seeman *et al* 2017). Similar to paired-pulse conditioning, spatiotemporal patterned stimulation can be used to apply stimulation across many sites with differing delays and amplitudes between sites. This type of patterned stimulation might be particularly advantageous for treating neuropathologies, such as stroke and Alzheimer’s, that result in aberrant network activity across multiple nodes. (Asp *et al* 2023; Ip *et al* 2021; Khateeb *et al* 2019; Khateeb *et al* 2022; Sato *et al* 2022; Wang *et al* 2013; Zhou *et al* 2022; Zhou *et al* 2023).

While paired-pulse and spatiotemporal stimulation are open-loop approaches, it is thought that a significant factor contributing to the inconsistent effects of neural stimulation is the variable brain states in which the stimulation is delivered (Bloch *et al* 2019; Bloch *et al* 2022; Zrenner and Ziemann 2024). Advancements in technology for rapidly processing ongoing neural activity have made it possible to deliver closed-loop stimulation during specific neural states. Providing support for cycle-triggered stimulation was motivated by several studies which found that delivering stimulation during a specific LFP phase resulted in larger stimulation evoked responses (Zanos *et al* 2018; Zrenner and Ziemann 2024; Zrenner *et al* 2018, Wischnewski *et al* 2022).

### 4.2. Comparison to other models

We chose to implement these features within the existing VERTEX software because unlike many other computational models that simulate spiking activity, LFPs, and synaptic plasticity in neurons, VERTEX uniquely does so in a large population of neurons using realistic biophysical properties. LFPy and The NEURON simulator are python-based models that predict spiking activity in highly realistic neuron models with more compartments and complex branching than VERTEX (Hines and Carnevale 1997; Lindén *et al* 2010). However, both are designed to simulate activity in a single neuron or a very small collection of neurons. In contrast, The Brian simulator uses point neurons but can simulate activity in a large population of neurons and has support for synaptic plasticity including STDP (Goodman and Brette 2009). Similarly, the integrate-and-fire model by Shupe and Fetz (2021) simulates point neurons without physical properties in several hundred neurons. It also incorporates STDP and various open- and closed-loop stimulation protocols. Despite advantageous features in other models, VERTEX’s use of realistic neuron morphologies and connectivity, where dendritic and synaptic activity contribute to LFPs, generates more realistic LFPs. This is particularly important as it allows users to explore the relationship between spikes and LFPs, an area with limited *in-vivo* research (Yazdan-Shahmorad *et al* 2011; Yazdan-Shahmorad *et al* 2013). By deepening our understanding of the correlations between spiking and LFPs, experimentalists could make greater use of LFP signals, which are obtained through less invasive methods.

### 4.3. Future directions

While our novel extensions provide comprehensive features to VERTEX, there is potential for further expansion and improvement of these simulations. In particular, our optogenetic stimulation model is based on several theoretical frameworks. More *in-vivo* research could refine these models to more accurately represent light spread through brain tissue, better account for light source parameters such as the optical fiber’s numerical aperture, and improve photocurrent dynamics for more realistic onset mechanics and longer duration light pulses to accommodate both peak and plateau currents.

## 5. Conclusions

Our extensions to VERTEX provide a highly adaptable, comprehensive, and realistic platform for users to test and predict the effects of diverse neural stimulation methods on spiking activity and local field potentials. We anticipate that these extensions will be highly valuable in the fields of systems neuroscience and therapeutic neural interfaces. These new features enable the exploration of numerous important questions, such as comparing the effects of optogenetic stimulation to electrical stimulation. At an individual level, for experimentalists, we hope these tools will serve as a preliminary means to predict local and network-level effects of modern stimulation methods before conducting *in-vivo* experiments. Doing so will reduce the number of extraneous hypotheses tested *in-vivo*, thereby saving costs, time, and reducing the use of animals.

## Acknowledgments

This work was funded by NIH R01NS119593, NSF 2223495, and the American Heart Association. We thank Riya Jain and Julien Bloch for their contributions to coding and parameterization.

## Code availability

All software needed to run model simulations and figure generation can be found at this code repository: https://github.com/lshupe/Vertex2_YL

## Conflict of interest

The authors declare no conflict of interest.

## Notes

https://github.com/lshupe/Vertex2_YL

### Competing Interest Statement

The authors have declared no competing interest.

### Summary of Updates

This version of the manuscript includes modeling the inhibitory opsin Jaws.

