## Supplemental Table 1 for "Flexible modeling of large-scale neural network stimulation: electrical and optical extensions to The Virtual Electrode Recording Tool for EXtracellular Potentials (VERTEX)"

| File name (.m) | Folder | Description |
|---|---|---|
| nsg_main_stdp | vertex\ | Main script for setting top-level parameters and activating STDP/stimulation protocols. This is sometimes saved to a more descriptive name to preserve a particular set up. |
| run_bsf_model_stdp_sim | \bsfModel_yazdan\ | Main function for running the simulation. Called from nsg_main_stdp.m (BSF stands for Brain Structure and Function, the journal which published the first VERTEX paper). |
| bsf_connectivity_dist | \bsfModel copy\ | Copy of original connectivity set up with options for randomizing the initial weights. |
| bsf_stdp | \bsfModel copy\ | Setup script for Spike-Time-Dependent Plasticity (STDP). |
| bsf_chr2_single_pulse | \bsfModel copy\ | Setup script for single pulse optogenetic stimulation using the ChR2 model. |
| bsf_vfChrimson_single_pulse | \bsfModel copy\ | Setup script for single pulse optogenetic stimulation using the vfChrimson model. |
| bsf_chronos_single_pulse | \bsfModel copy\ | Setup script for single pulse optogenetic stimulation using the Chronos model. |
| bsf_jaws_single_pulse | \bsfModel copy\ | Setup script for single pulse optogenetic stimulation using the Jaws model. |
| bsf_chr2_pattern | \bsfModel copy\ | Setup script for multi-pulse pattern optogenetic stimulation using the ChR2 model. |
| bsf_surface_single_pulse | \bsfModel copy\ | Setup script for single pulse field stimulation using surface electrodes. |
| bsf_stim_single_pulse | \bsfModel copy\ | Setup script for single pulse field stimulation in Layer2/3. |
| bsf_stim_paired_pulse | \bsfModel copy\ | Setup script for paired pulse field stimulation. |
| bsf_stim_pattern | \bsfModel copy\ | Setup script for multi-pulse patterned field stimulation. |
| BipolarElectrodeSurfaceGeometry | \vertex_stimulation\ | Function to create patch electrode geometry (e.g. |

| | | |
|---|---|---|
| | | electrodes on the tissue surface.) |
| BipolarElectrodeVerticalGeometry | \vertex_stimulation\ | Function to create tip electrode geometry (e.g. electrodes penetrating from the tissue surface.) |
| EstimateElectrodeCurrent | \vertex_stimulation\ | Example Function for estimating current delivered at at tip electrode. |
| runSimulation_stim | \vertex_user\ | Modified version of the original runSimulation.m function. Called from run_bsf_model_stdp. |
| simulateParallel_stim | \vertex_simulate\parallel\ | Modified version of the original simulateParallel.m function. Called from runSimulaton_stim. |
| opto_current_chr2 | \opto\ | ChR2 irradiance to photocurrent function. |
| opto_current_vfChrimson | \opto\ | vfChrimson irradiance to photocurrent function. |
| opto_current_chronos | \opto\ | Chronos irradiance to photocurrent function. |
| opto_current_jaws | \opto\ | Jaws irradiance to photocurrent function. |
| opto_irradiance_blue_473nm | \opto\ | Function to model blue light spread through tissue. |
| opto_irradiance_yellow_594nm | \opto\ | Function to model yellow (amber) light spread through tissue. |
| InputModel_i_opto | \vertex_simulate\models\inputs\ | VERTEX Input Model for optogenetic photocurrent. |
| testfigs | vertex\ | Function for creating summary plots given a path to a simulation's results folder. |
| compare_weights | \model_analysis\ | Function for comparing synaptic weights saved at two different times during the same simulation. |

**Supplemental Table 1: New code modules.** Code modules that were modified from the original VERTEX codebase, or were added to make our described extensions.
