## Supplemental Figure 1 for "Flexible modeling of large-scale neural network stimulation: electrical and optical extensions to The Virtual Electrode Recording Tool for EXtracellular Potentials (VERTEX)"

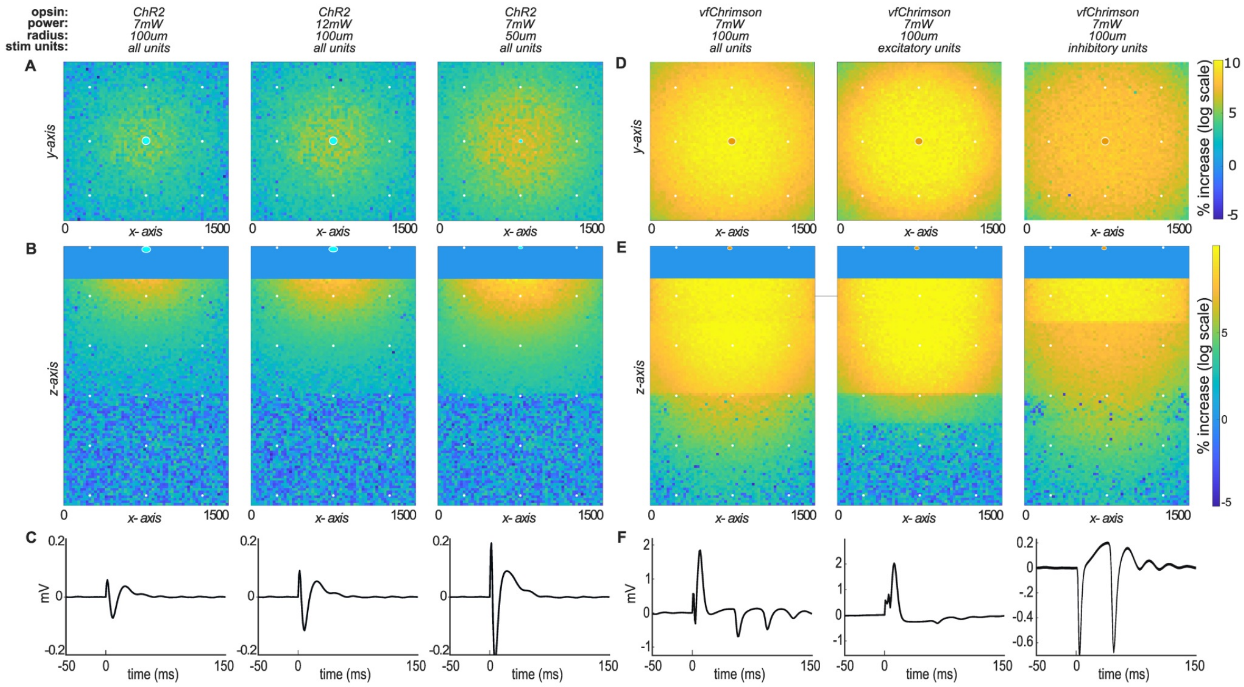


**Supplemental Figure 1. Simulations using various modifiable optogenetic parameters.** A) Top-down and B) side-view of change in spiking activity after optogenetic stimulation using the ChR2/blue light model under different light powers or light source radii. C) LFPs at the surface-center recording electrode for each simulation. D) Top-down and E) side-view of change in spiking activity using the vfChrimson/amber light models where different neuron unit groups/cell types are light-responsive for each of the three simulations. In the first column all neurons are light-responsive, whereas only excitatory or inhibitory neurons are set as light-responsive in the second and third column, respectively. F) LFPs at the surface-center recording electrode for each simulation.
